# Decoding the silence: Neural bases of zero pronoun resolution in Chinese

**DOI:** 10.1101/2021.05.06.442989

**Authors:** Shulin Zhang, Jixing Li, Yiming Yang, John Hale

## Abstract

Chinese is one of many languages that can drop subjects. We report an fMRI study of language comprehension processes in these “zero pronoun” cases. The fMRI data come from Chinese speakers who listened to an audiobook. We conducted both univariate GLM and multivariate pattern analysis (MVPA) on these data time-locked to each verb with a zero pronoun subject. We found increased left middle temporal gyrus activity for zero pronouns compared to overt subjects, suggesting additional effort searching for an antecedent during zero pronoun resolution. MVPA further revealed that the intended referent of a zero pronoun seems to be physically represented in the Precuneus and the Parahippocampal Gyrus shortly after its presentation. This highlights the role of memory and discourse-level processing in resolving referential expressions, including unspoken ones, in naturalistic language comprehension.

## 1 Introduction

Our ability to ascertain which entity a pronoun refers to is a central part of human language understanding. Many East Asian languages, such as Mandarin Chinese, pronouns can be freely omitted in both subject and object positions given proper discourse context (Huang, 1989). This phenomenon of “zero pronoun resolution” has been extensively studied in formal linguistics (e.g., Barbosa, 2011, 2019; Bi & Jenks, 2019; C. N. Li & Thompson, 1976; Neeleman & Szendrői, 2007; Song, 2005), yet its neural bases are barely discussed, especially with naturalistic stimuli that can reveal language processes at the discourse level. Here we report the results of the first fMRI study to examine the brain regions involved in zero pronoun processing while Chinese participants listen to a naturalistic narrative.

The status of zero pronouns as the deleted counterparts of regular pronouns is debated in formal linguistics. While some assumed that null pronouns are overt pronouns that fail to be realized at the phonological interface (e.g., Neeleman & Szendrői, 2007), others suggested that null pronouns are derived from semantically distinct noun phrases (Bi & Jenks, 2019). From a cognitive perspective, zero pronouns are the “missing spots” in texts and speech, and they constitute a “harder” case for pronoun resolution as they offer no phonological or morpho-syntactic information. By comparing brain activity during the processing of zero and non-zero arguments, we aim to better understand the neural mechanisms involved in understanding unpronounced pronouns in *pro*-drop languages.

### 1.1 Brain regions involved in reference processing

While no prior neuroimaging study has directly investigated zero pronoun processing, there are some fMRI and MEG studies on referential processing in general (Brodbeck & Pylkkänen, 2017; Brodbeck et al., 2016; Hammer et al., 2007; J. Li et al., 2021; Matchin et al., 2014; Nieuwland et al., 2007; Santi & Grodzinsky, 2012). However, no consensus has been reached on the neural correlates for pronoun processing. In addition, previous studies adopted different task manipulations, making it unclear whether they tapped the same cognitive processes. For example, Nieuwland et al. (2007) compared the BOLD responses when participants read sentences containing a “referentially failing pronoun” (*e*.*g*., “Rose told Emily that *he* had a positive attitude towards life.”) or a coherent pronoun (e.g., “Ronald told Emily that *he* had a positive attitude towards life.”). Nieuwland et al. showed that referentially failing pronouns were associated with increased activation in the medial parietal regions and bilateral inferior parietal regions, possibly reflecting morpho-syntactic processing. Hammer et al. (2007) manipulated the syntactic gender matching between the antecedent and pronouns using German sentences and found that gender incongruency elicited the bilateral Inferior Frontal Gyrus (IFG), the left Medial Frontal Gyrus (MFG), and the bilateral Supramarginal/Angular Gyrus compared to congruent pronoun-antecedent pairs. Hammer et al. (2011) further investigated the possible interactions between gender and distance between the antecedents and the pronoun. The results showed a fronto-temporal network including the bilateral IFG, the Superior Temporal Gyrus (STG), and posterior Middle Temporal Gyrus (pMTG) for long-distance conditions, with the pMTG additionally driven by syntactic gender violation. These authors suggested that the temporal regions are sensitive to the morpho-syntactic information of the antecedents since the long distance between the antecedent and the pronoun increased the overall syntactic complexity of the sentence. Matchin et al. (2014) also examined the effect of distance but with the backward anaphora/filler-gap dependencies contrast. Matchin and colleagues observed specific activity in the bilateral Anterior Temporal Lobes (ATLs), the bilateral Angular Gyrus (AGs), and the left Precuneus activity during the processing of backward anaphora compared to *wh*-fillers. Santi & Grodzinsky (2012) compared null pronouns, a parasitic-gap and a *wh*-trace in English sentences such as “[Which paper] did the tired student submit [*wh*-trace] after reviewing [parasitic gap/it]?”. The results showed increased activity in the right Middle Frontal Gyrus (MFG), the left Ventral Precentral Sulcus, and the Left Supramarginal Gyrus for pronouns compared to parasitic gaps.

In addition to the morpho-syntactic manipulations, Brodbeck & Pylkkänen (2017) and Brodbeck et al. (2016) used a visual world paradigm in magnetoencephalography (MEG) and found medial parietal activity in cases of successful reference resolution. More relevant to the current study is J. Li et al.’s (2021) study on third person pronoun processing using the same naturalistic listening paradigm. In both fMRI and MEG, Li et al. found that the left middle temporal gyrus (LMTG) is consistently activated for third person pronoun processing in both English and Chinese. Yet they also found additional medial parietal activity from the MEG data, consistent with Brodbeck & Pylkkänen (2017), Brodbeck et al. (2016), and Nieuwland et al. (2007).

To sum up, referential processing has been implicated in a number of regions, including the medial parietal lobe. Zero pronoun resolution, as a special case of referential processing, is expected to involve similar brain regions.

### 1.2 Zero pronouns in Chinese

As a “radical *pro*-drop” language, Chinese can have a null pronoun as the subject or object of a tense clause in appropriate contexts. Unlike ordinary “*pro*-drop” languages, such as Spanish and Italian, that exhibit rich verbal agreement systems, Chinese does not have verbal inflections that provide person or gender information to help recover the omitted pronouns (See (1), data from Huang (1989)). Instead, in Chinese, zero pronouns and their overt coreferential noun phrases form a topic chain structure, a discourse structure that enables covert as well as overt coreference (Kun, 2019; W. Li, 2004; Shi, 1993).

1. “Zhangsan kanjian Lisi le ma?” (“Did Zhangsan see Lisi?”)
  a. “Ta kanjian ta le.” (“He saw him.”)
  b. “[] kanjian ta le.” (“[He] saw him.”)
  c. “Ta kanjian [] le.” (“He saw [him].”)
  d. “[] kanjian [] le.” (“[He] saw [him].”)

A topic chain is a chain of clauses sharing an identical topic that occurs overtly once in one of the clauses, and its boundary may cross several sentences and even paragraphs (W. Li, 2004). The topic chain can integrate information from multiple clauses (Kun, 2019), which makes long-distance coreference between zero pronouns and overt noun phrases possible. We can understand coreference resolution as searching for an appropriate antecedent in the topic chain. This searching process likely recruits memory and discourse-related brain regions.

The existence of the topic chain, coupled with the lack of morphological markers, makes zero pronoun resolution a “harder” case of pronoun resolution in Chinese, such that additional cognitive resources may be needed to recover the omitted arguments. We would expect the involvement of brain regions related to discourse-level processing in Chinese zero pronoun resolution, and higher brain activation level compared with overt noun phrases.

### 1.3 Current study

The current study examines which brain regions are responsible for the processing of the dropped pronouns in Chinese. We compared brain activity time-locked to zero and non-zero subjects during naturalistic listening. Since zero pronouns are not pronounced in the speech, we marked the onsets of the main verbs that follow either a zero or an overt subject as the time point where the zero/non-zero argument occurs (See Section 2.2 for details on the annotation steps).

In a mass univariate analysis with a General Linear Model (GLM), we show that zero pronoun resolution demands higher activity in anterior as well as posterior LMTG, compared to overt reference resolution (See Section 3.1). Given the LMTG’s role in pronoun resolution (Hammer et al., 2007, 2011; J. Li et al., 2021, e.g.,), our results suggest that zero pronoun resolution evokes additional effort expended in the search for an antecedent. With searchlight-based Multivariate Pattern Analysis (MVPA), we identify a network that includes the Precuneus and Parahippocampal Gyrus, which are regarded as part of the “extended” language network, compared to the “core” language network including brain regions such as the temporal lobe (Fedorenko et al., 2011; Ferstl et al., 2008; Xiong & Newman, 2021). These results suggest that brain regions beyond the “core” language network subserve zero pronoun resolution in Chinese.

## 2 Material and methods

### 2.1 Participants

Participants were 35 healthy, right-handed, young adults (15 female, mean age=19.3, range = 18-25). They self-identified as native Chinese speakers and had no history of psychiatric, neurological, or other medical illness that could compromise cognitive functions. All participants were paid for and gave written informed consent prior to participation, in accordance with the guidelines of the Ethics Committee at Jiangsu Normal University.

### 2.2 Stimuli and annotations

The stimuli is a Chinese translation (xiaowangzi.org, 2021) of Saint-Exupéry’s *The Little Prince*. To annotate zero and non-zero subjects, we first located all verbs (i.e., “VV”s) in the text using ZPar (Zhang & Clark, 2011). We then annotated each verb as Zero or Non-zero based on whether it has an overtly pronounced subject. For example, as shown in Figure 1a, the verb phrase (VP) “弄不懂 (make no understanding)” is marked as non-zero as its subject “大人们 (grown-ups)” is overt; the VP “做解释 (make explanations)” is marked as “zero” since its subject “我 (I)” is omitted.

**Figure 1:**
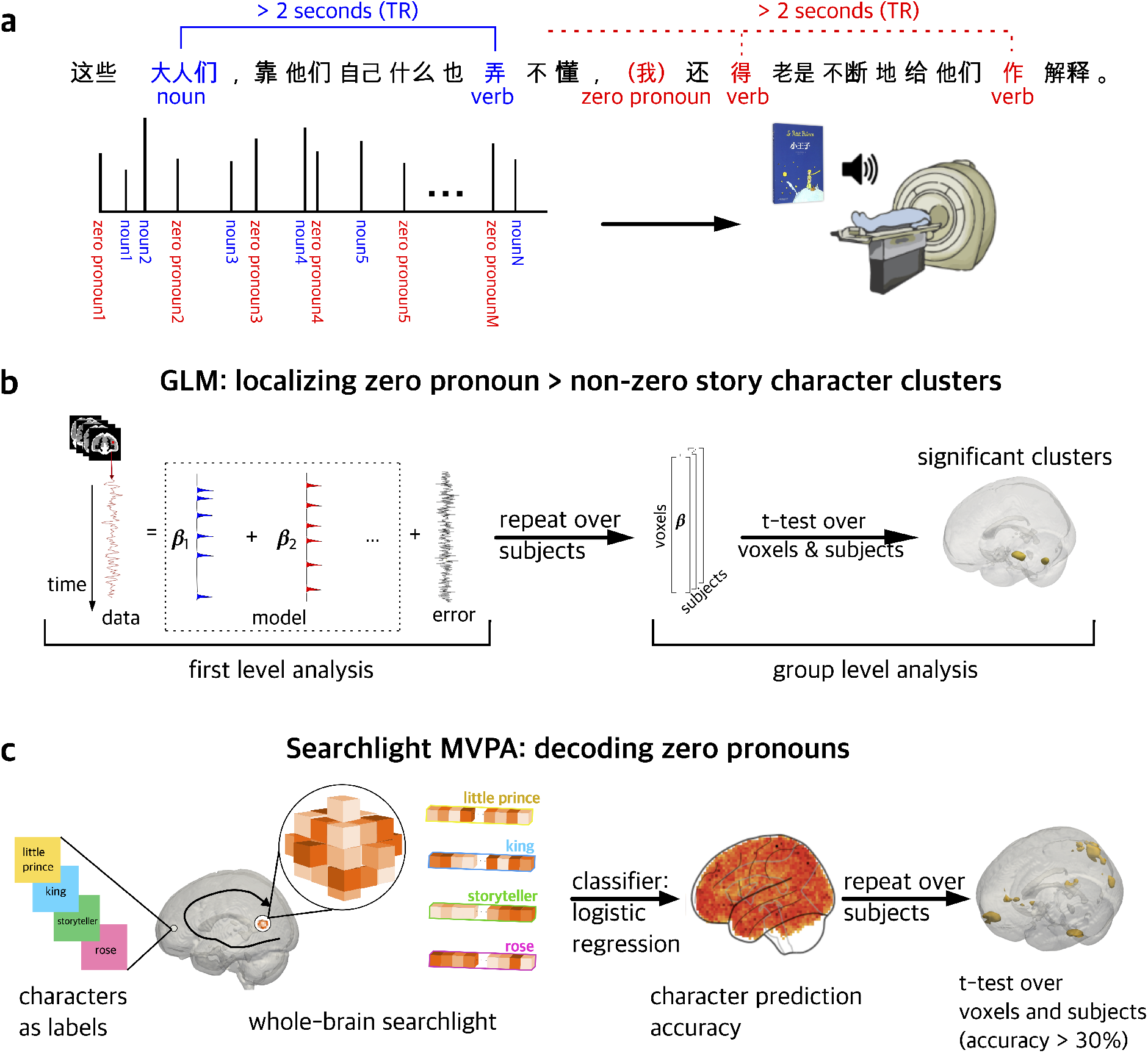
Schematic illustration of the analysis procedure. **a**. Stimuli and fMRI data collection. fMRI data were collected while Chinese native speakers were listening to a naturalistic audiobook. Zero pronouns and Non-zero nouns in subject position are annotated by Chinese native speakers, and their corresponding main verbs were taken as timestamps for GLM and MVPA analyses. The distance between a zero or non-zero noun to its main verb was controlled to be longer than 2 seconds so that they cannot be in the same fMRI scan. **b**. Two-stage General Linear Model analyses. At the first stage, a general linear model was fitted to each participants’ fMRI data, and the regressors used in the model include audio sound pressure, word frequency, zero-pronoun feature, non-zero noun feature (See Section 2.5). At the second stage, a *t*-test was performed on the distribution of *β* values across subjects and voxels, and the significant clusters were retrieved with *p <* .05 FWE and *k >* 20. **c**. Whole-brain searchlight multivariate pattern analysis. Among all annotated zero pronouns, four story characters’ (See Section 2.6) main verb scans were used in the MVPA decoding analyses. A logistic regression classifier was used to derive an average accuracy value for decodability of the four story characters, based on an N-voxel neighborhood. T-tests were performed on these accuracy values across subjects to identify clusters with above-chance accuracy. (*p <* .001 FWE and *k >* 50).

### 2.3 Procedure

After giving their informed consent, participants were familiarized with the MRI facility and assumed a supine position on the scanner. The presentation script was written in PsychoPy 2 (Peirce, 2007). Auditory stimuli were delivered through MRI-safe, high-fidelity headphones (Ear Bud Headset, Resonance Technology, Inc, California, USA) inside the head coil. The headphones were secured against the plastic frame of the coil using foam blocks. An experimenter increased the sound volume stepwise until the participants could hear clearly.

The Chinese audiobook lasted for about 99 minutes and was divided into nine sections, each lasted for about ten minutes. Participants listened passively to the nine sections and completed four quiz questions after each section (36 questions in total). These questions were used to confirm their comprehension and were viewed by the participants via a mirror attached to the head coil and they answered through a button box. The entire session lasted for around 2.5 hours.

### 2.4 fMRI data collection and preprocessing

MRI images were acquired with a 3T MRI GE Discovery MR750 scanner with a 32-channel head coil. Anatomical scans were acquired using a T1-weighted volumetric Magnetization Prepared RApid Gradient-Echo (MP-RAGE) pulse sequence. Functional scans were acquired using a multi-echo planar imaging (ME-EPI) sequence with online reconstruction (TR=2000 ms; TEs=12.8, 27.5, 43 ms; FA=77°; matrix size=72 x 72; FOV=240.0 mm x 240.0 mm; 2 x image acceleration; 33 axial slices, voxel size=3.75 x 3.75 x 3.8 mm). Cushions and clamps were used to minimize head movement during scanning.

All fMRI data were preprocessed using AFNI version 16 (Cox, 1996). The first 4 volumes in each run were excluded from analyses to allow for T1-equilibration effects. Multi-echo independent components analysis (ME-ICA) (Kundu et al., 2012) was used to denoise data for motion, physiology, and scanner artifacts. Images were then spatially normalized to the standard space of the Montreal Neurological Institute (MNI) atlas, yielding a volumetric time series resampled at 2 mm cubic voxels.

### 2.5 GLM analysis

A whole-brain GLM analysis was conducted to localize the brain regions involved in zero and non-zero reference resolution. We modeled the timecourse of each voxel’s BOLD signals for each of the nine sections by a binary zero pronoun regressor and a binary non-zero subject regressor, time-locked to the onset of the verb for the zero subject (510 cases) and non-zero subject (1942 cases) in the audiobook. We included three control variables: the root mean square intensity (*RMS intensity*) for every 10 ms of each audio section, the binary regressor time-locked to the offset of each word in the audio (*word rate*), and the unigram frequency of each word (*frequency*), estimated using Google ngrams (Version 20120701) and the SUBTLEX corpora for Chinese (Cai & Brysbaert, 2010). These regressors were convolved with SPM12’s (Penny et al., 2011) canonical HRF function and matched the scan numbers of each section. (See Supplementary Figure 5 for the correlation matrix of the regressors, and Supplementary Figure 4 for a visualization of the regressors.) At the group level, the contrast images for zero and non-zero subjects were examined by a factorial design matrix. An 8 mm full-width at half-maximum (FWHM) Gaussian smoothing kernel was applied on the contrast images from the first-level analysis to counteract inter-subject anatomical variation. Significant clusters were thresholded at *p <* .05 FWE with cluster size of *k >* 20. The GLM analysis was performed with the python package nilearn (0.7.0) (Abraham et al., 2014).

### 2.6 MVPA for zero pronoun resolution

A whole-brain searchlight MVPA was performed to discriminate patterns of activation pertaining to the omitted story characters. The fMRI scans which contain both a zero pronoun and its previous overtly pronounced antecedent are excluded from the MVPA. We selected the four most frequent story characters for the classification, and there were 188 zero-pronoun instances used in MVPA, including: “小王子(the little prince)”, 84 instances; “我(I/the storyteller)”, 67 instances; “国王 (the king)”, 25 instances; “花 (the rose)”, 12 instances.

Searchlight MVPA identifies voxels where the pattern of activation in its local neighborhood can discriminate between conditions (i.e. story characters). For each subject, a spherical ROI (radius = 8 mm) centered in turn on each voxel in the brain scans time-locked to 5 seconds after the zero pronouns’ presentation. A 5-second delay serves to capture BOLD signals at approximately the peak of their hemodynamic response to the zero pronouns. Each vector contains all the voxels in each sphere without feature selection. A logistic regression classifier was trained to differentiate the vectors of all four story characters. A 3-fold cross-validation process was adopted in the training process, which means 2/3 of the original labeled data were used as a training dataset, and the rest as a testing set. Prediction accuracy was averaged over the three testing results.

This whole process was repeated for the sphere centered by each voxel for each subject. The resulting maps contain each voxel’s decoding accuracy for each subject. Higher accuracy indicates better performance on decoding the reference of the zero pronouns. At the group level, a *t*-test was conducted for all voxels across all subjects. Voxels with an accuracy higher than 30% (higher than the chance baseline 25% ^1^) were highlighted. Family-wise error correction was applied with an alpha level of *<*.001 and an adequate cluster size of *k >* 50. The MVPA analysis was performed using the python packages nilearn (0.7.0) (Abraham et al., 2014), and scikit-learn (Pedregosa et al., 2011).

To characterize chance performance, we carried out another MVPA analysis on a scrambled dataset where story character labels were assigned randomly. In this supplementary analysis, a story character label randomly selected out of the four story characters was assigned to each zero pronoun, and the same MVPA analysis steps introduced above were conducted to test whether there are brain regions able to decode randomly assigned labels.

## 3 Results

### 3.1 GLM: Localizing brain regions for zero pronoun processing

The contrast between zero pronouns and overt references to story characters revealed significantly higher activity in the anterior and posterior LMTGs (*p <* .001 FWE, peak *t*-value = 6.02, cluster size = 680 mm^3^ and *p <* .001 FWE, peak *t*-value = 5.87, cluster size = 1344 mm^3^, respectively; see Figure 2a,b). The MNI coordinates for the peak of each cluster are shown in Figure 2c. No significant cluster was found for the opposite contrast, i.e. Non-zero *>* Zero.

**Figure 2:**
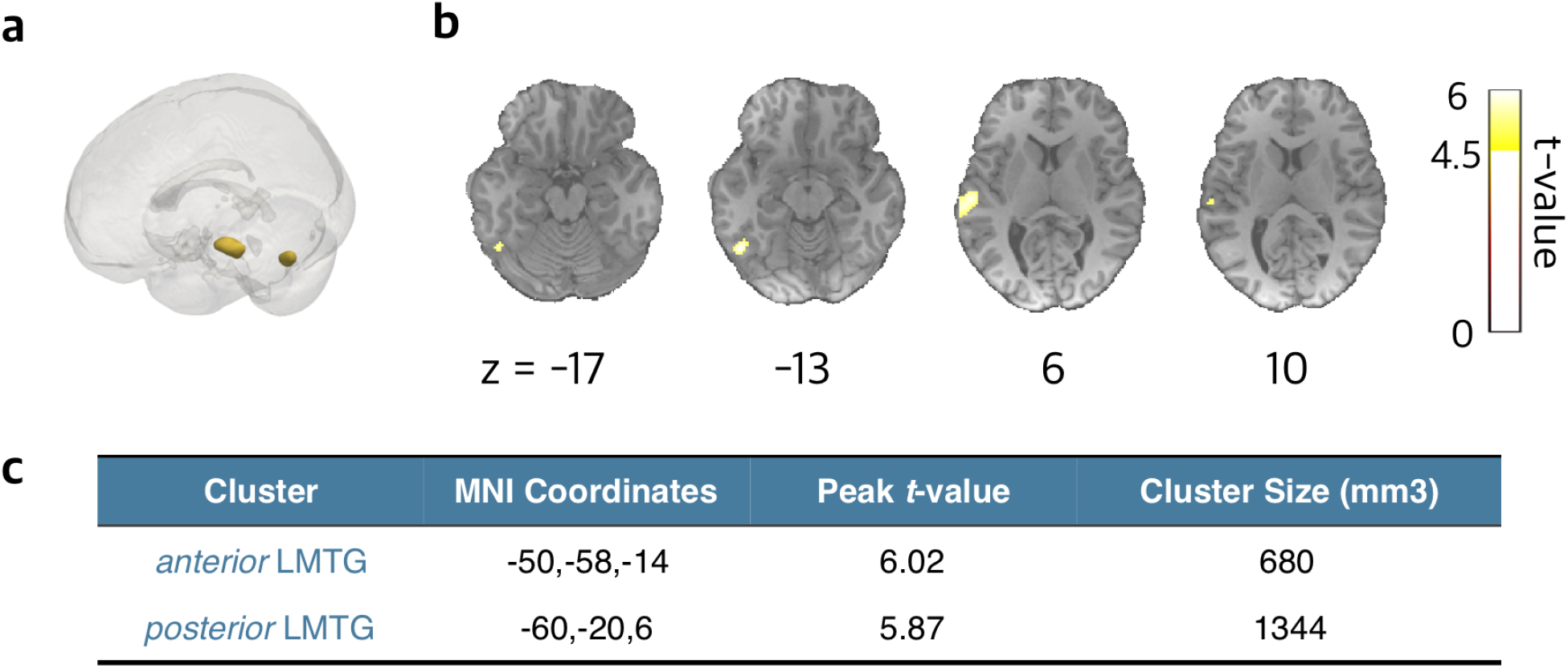
GLM results for the contrast between zero and non-zero reference resolution. **a** Whole-brain view on a 3D brain. **b** Coronal slices of significant clusters **c** MNI coordinates, cluster size and their peak level statistics, thresholded at *p <* .05 FWE and *k >* 20.

### 3.2 MVPA: Decoding references of zero pronouns

Searchlight MVPA results are shown in Figure 3. Brain regions with a decoding accuracy greater than 30% for the zero pronouns include the Precuneus (the right Precuneus: *p <* .001 FWE, peak *t*-value = 12.9, cluster size = 3027 mm^3^; the left Precuneus: *p <* .001 FWE, peak *t*-value = 11.78, cluster size = 5676 mm^3^), the LMFG (*p <* .001 FWE, peak *t*-value = 11.76, cluster size = 1608 mm^3^), the right Interior Temporal Gyrus (RITG; *p <* .001 FWE, peak *t*-value = 11.15, cluster size = 2176 mm^3^), the LAG (*p <* .001 FWE, peak *t*-value = 10.78, cluster size = 1955 mm^3^), the LMTG (*p <* .001 FWE, peak *t*-value = 10.55, cluster size = 2522 mm^3^), the left Frontal Pole (*p <* .001 FWE, peak *t*-value = 10.29, cluster size = 2680 mm^3^), the RAG (*p <* .001 FWE, peak *t*-value = 10.01, cluster size = 1671 mm^3^), and the right Parahippocampal Gyrus (*p <* .001 FWE, peak *t*-value = 9.27, cluster size = 2712 mm^3^).

**Figure 3:**
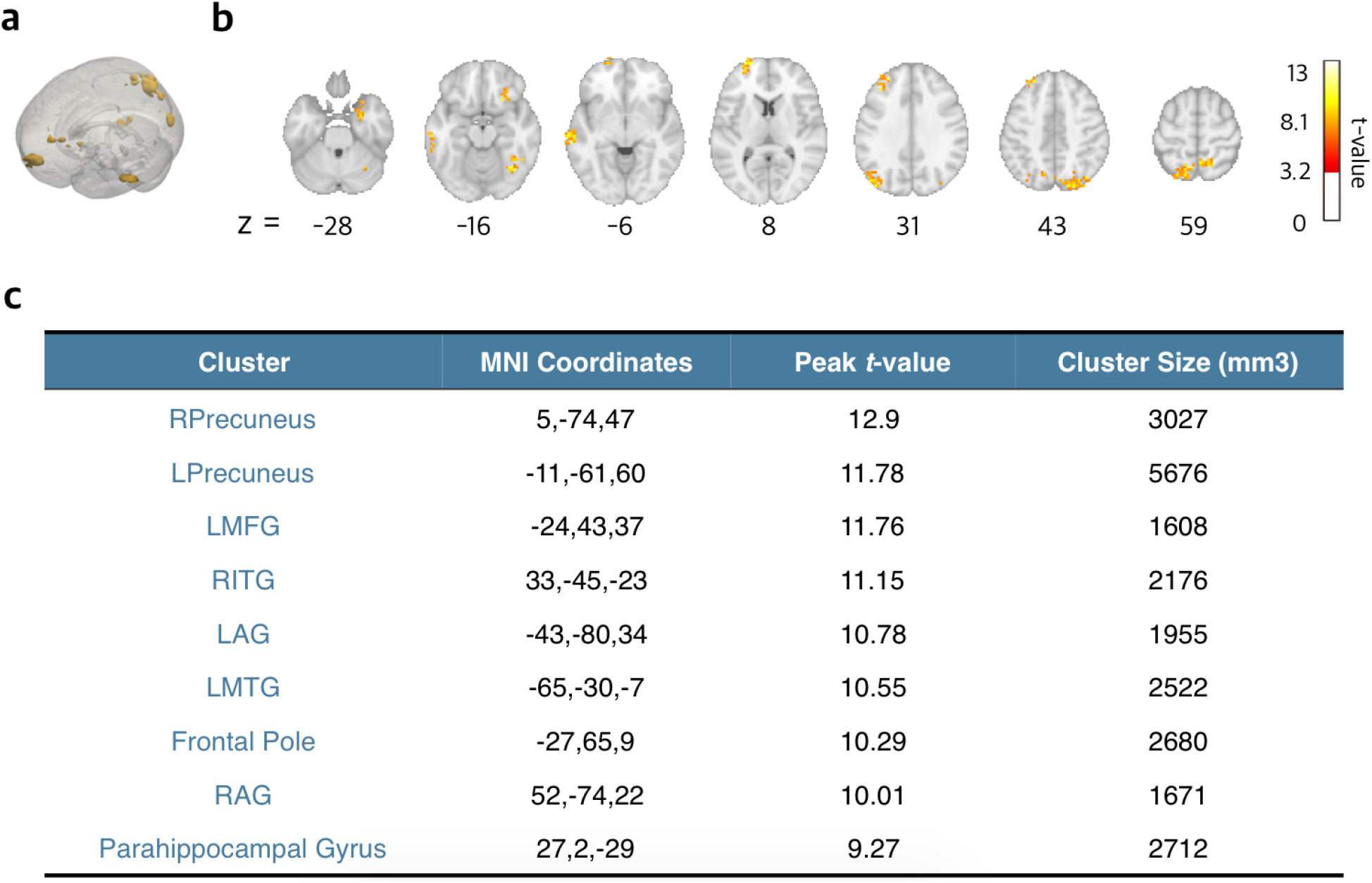
MVPA results for brain regions with decoding accuracy significantly higher than 30%. **a** Whole-brain view on a 3D brain. **b** Coronal slices of significant clusters **c** MNI coordinates, cluster size and their peak level statistics, thresholded at *p <* .001 FWE and *k >* 50.

Both the GLM and MVPA results implicate the LMTG. Not only did the LMTG show higher activity for zero pronoun resolution compared to non-zero reference resolution, but also showed high story character decoding accuracy for story characters. MVPA further revealed a network for decoding zero pronouns, including the Precuneus, the LAG, the frontal pole, the LMFG, and the Parahippocampal Gyrus.

### 3.3 Scrambled MVPA: Decoding zero pronouns with randomly assigned story character labels

When the correct story character labels are replaced by randomly assigned story character labels, no significant brain region was detected from the MVPA results (*p <* .001 FWE, cluster size *>* 50). The null result with randomly-assigned label supports the idea that the main MVPA analysis in Section 3.2 in fact does identify brain regions where story character information is represented.

## 4 Discussion

This study examines the neural bases of zero pronoun resolution in Chinese. Chinese is especially suitable for studying zero pronoun resolution as it does not have verbal inflections that could interfere with zero pronoun resolution at the phonological and morpho-syntactic levels. The GLM results show increased LMTG activity during zero pronoun resolution compared to non-zero reference processing. MVPA results further reveal a network of activity including the Precuneus and the Parahippocampal Gyrus in addition to the “core” language network. The results suggest that zero pronoun resolution involves additional effort in the search of an antecedent compared to regular noun phrases. Both “core” and “extended” nodes of the language network appear to contribute to resolving the reference of zero pronouns.

### 4.1 LMTG for retrieving the antecedents during zero pronoun resolution

Both anterior and posterior regions within the LMTG showed significantly higher activity for zero pronouns compared to overt references to story characters. In previous studies, the LMTG has been shown to play an essential role in language comprehension (Dronkers et al., 2011; Matchin & Hickok, 2020). The LMTG has also been associated with biological and syntactic gender processing (Heim et al., 2002; Hammer et al., 2007, 2011; Miceli et al., 2002) during pronoun processing. For example, Hammer et al. (2007) showed that German sentences with congruent biological and syntactic gender evoked higher activity in the LMTG; Miceli et al. (2002) found increased LMTG activity when the subjects were asked whether a written noun has a masculine or feminine gender. However, J. Li et al. (2021) using the same naturalistic paradigm in fMRI, showed that the LMTG is also implicated for pronoun processing in Chinese. In addition, P. Li et al. (2004) showed a number of brain regions including the LMTG during a lexical-judgement task while Chinese participants saw nouns, verbs, and noun/verb-ambiguous words, supporting the LMTG’s role in lexical representation. Ferstl et al.’s (2008) meta-analysis for the language network further suggests that the LMTG contributes to the comprehension of coherence, and shows stable significant results as part of the “core” language network.

In the context of this existing evidence regarding LMTG’s role in morpho-syntactic matching and discourse coherence, increased LMTG activity for zero pronouns in the current study could reflect the greater difficulty of a reference-resolution problem in zero pronoun cases that lack phonological as well as morpho-syntactic information.

### 4.2 The neural network for zero pronoun resolution

Whole-brain searchlight-based MVPA revealed a network of brain regions implicated in the comprehension of reference to story characters, including the bilateral Precuneus, the bilateral AG, the left Frontal Pole, the LMFG, the LMTG, the RITG, and the right Parahippocampal Gyrus (See Figure 3).

The Precuneus has been previously related to “extra-linguistic” processing such as discourse-level information integration and memory retrieval (Bhattasali et al., 2019; Diachek et al., 2020; Foudil et al., 2020; Mashal et al., 2014; Wehbe et al., 2020). Foudil et al. (2020), for example, showed that brain activation level in the Precuneus was modulated by storyline consistency, suggesting its role in discourse information integration. (Mashal et al., 2014) found that schizophrenia patients with impaired capability towards metaphor comprehension showed higher activity in the left Precuneus compared to healthy participants. Bhattasali et al. (2019) using a same naturalistic listening paradigm, showed that the right Precuneus was correlated with the retrieval of stored expressions. On the other hand, the Precuneus is also suggested to be a “processing core” that connects to the MTG and the AG and integrates multiple brain functions such as memory retrieval (Mar, 2011). Here the results show that the Precuneus represents story character information, and this is consistent with the idea that the Precuneus is crucial for discourse-level processing.

The Parahippocampal Gyrus has also been implicated in discourse-level language processing (Allendorfer et al., 2012; Wallentin et al., 2005). For example, Allendorfer et al. (2012) showed higher Parahippocampal Gyrus activity while participants were generating verbs silently for a given noun. Wallentin et al. (2005) found the right Parahippocampal Gyrus activity while processing real motion sentences (*e*.*g*. “the man goes through the house”) and fictive motion sentences (*e*.*g*. “the trail goes through the house”). The decodability of story characters in the right Parahippocampal Gyrus further suggests that the search for antecedent involves discourse-level language processing. The Parahippocampal Gyrus, in previous studies, has also been reported relative to semantic memory retrieval and semantic verbal memory processing (Bartha et al., 2003). In Bartha et al.’s fMRI study, the subjects performed a semantic decision task while they heard spoken concrete nouns designating objects and made a decision on whether these objects were available in the supermarket and their costs compared to certain amounts. Bartha et al. observed activation in the Parahippocampal Gyrus, along with the medial temporal lobe and the inferior temporal lobe, and they inferred these brain regions’ relativity to semantic verbal memory processing. These results support the Parahippocampal Gyrus’s role for semantic language processing and discourse-level language processing as a brain region in the extended language network.

Apart from the Precuneus and the Parahippocampal Gyrus, we also identified a number of regions within the language network. The left AG has been suggested to support multimodal and multi-sensory associations that connect with brain regions for attention, episodic and semantic memory, and sentence level comprehension (Bonner et al., 2013; Humphreys et al., 2021; Price et al., 2015; Ramanan et al., 2018; Seghier, 2013). Using Transcranial Magnetic Stimulation (TMS), Branzi et al. (2021) found that the left AG is critical for integrating context-dependent information during language processing. Moreover, Davis & Yee (2019) suggested that the left AG’s connectivity to hippocampal regions underpins its essential role in processing thematic relations. The Frontal Pole is part of the deep track ventral pathway in the language network (Brauer et al., 2013) and is implicated for higher-level cognition processes, such as reasoning, episodic memory, and prospective memory (Tsujimoto et al., 2011). The left MFG is related to attention, working memory, and language processing (Briggs et al., 2021; Hazem et al., 2021). In a meta-analysis of fMRI studies by Wu et al. (2012), the LMFG had been found relevant for phonological and semantic processing in Chinese. The RITG has also been associated with language tasks such as metaphor and humor understanding (Ahrens et al., 2007; Bartolo et al., 2006) and noun processing (Crepaldi et al., 2013). To summarize, the brain network for resolving zero pronouns includes both the core language network and the extended language network e.g. the Precuneus and the Parahippocampal Gyrus. The involvement of brain regions related to discourse-level language processing and memory retrieval supports our previous assumption that zero pronoun resolution requires the involvement of these brain regions. Incidentally, these brain regions have been found to be closely connected under resting state functional connectivity analysis (Xu et al., 2019, 2015).

## 5 Conclusions

This study examines the neural bases of zero pronoun processing in Chinese. By comparing fMRI BOLD responses for zero pronoun processing with that of non-zero reference processing during naturalistic listening, we show that zero pronoun resolution evokes increased activity in the LMTG, suggesting additional effort in the search for an antecedent. By decoding brain activity patterns for zero pronouns with different references, we show a network of activity, including the Precuneus and the Parahippocampal Gyrus that are outside the core language network.

## A Supplemental Material

**Supplementary Figure 4:**
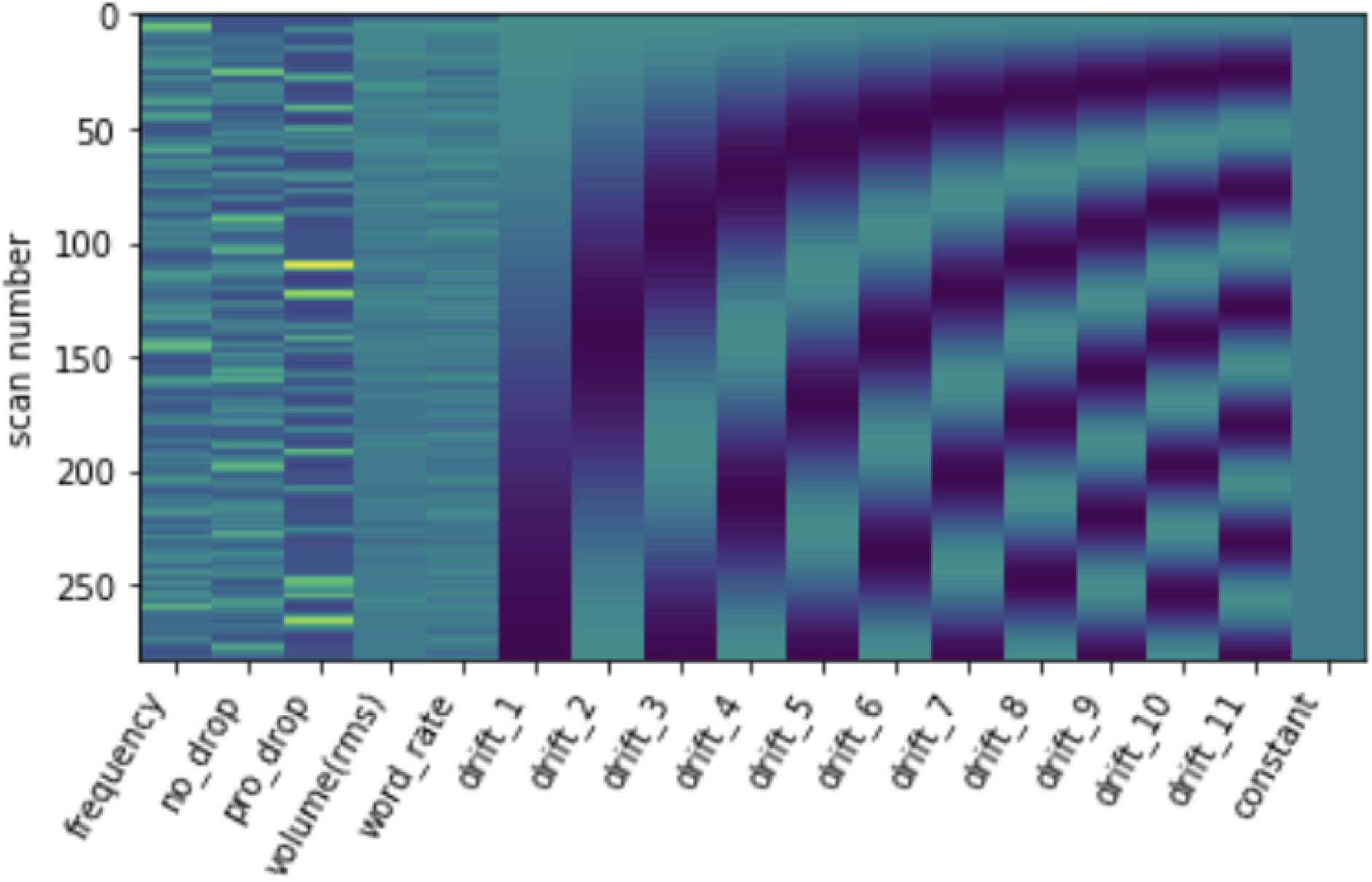
GLM analysis design matrix

**Supplementary Figure 5:**
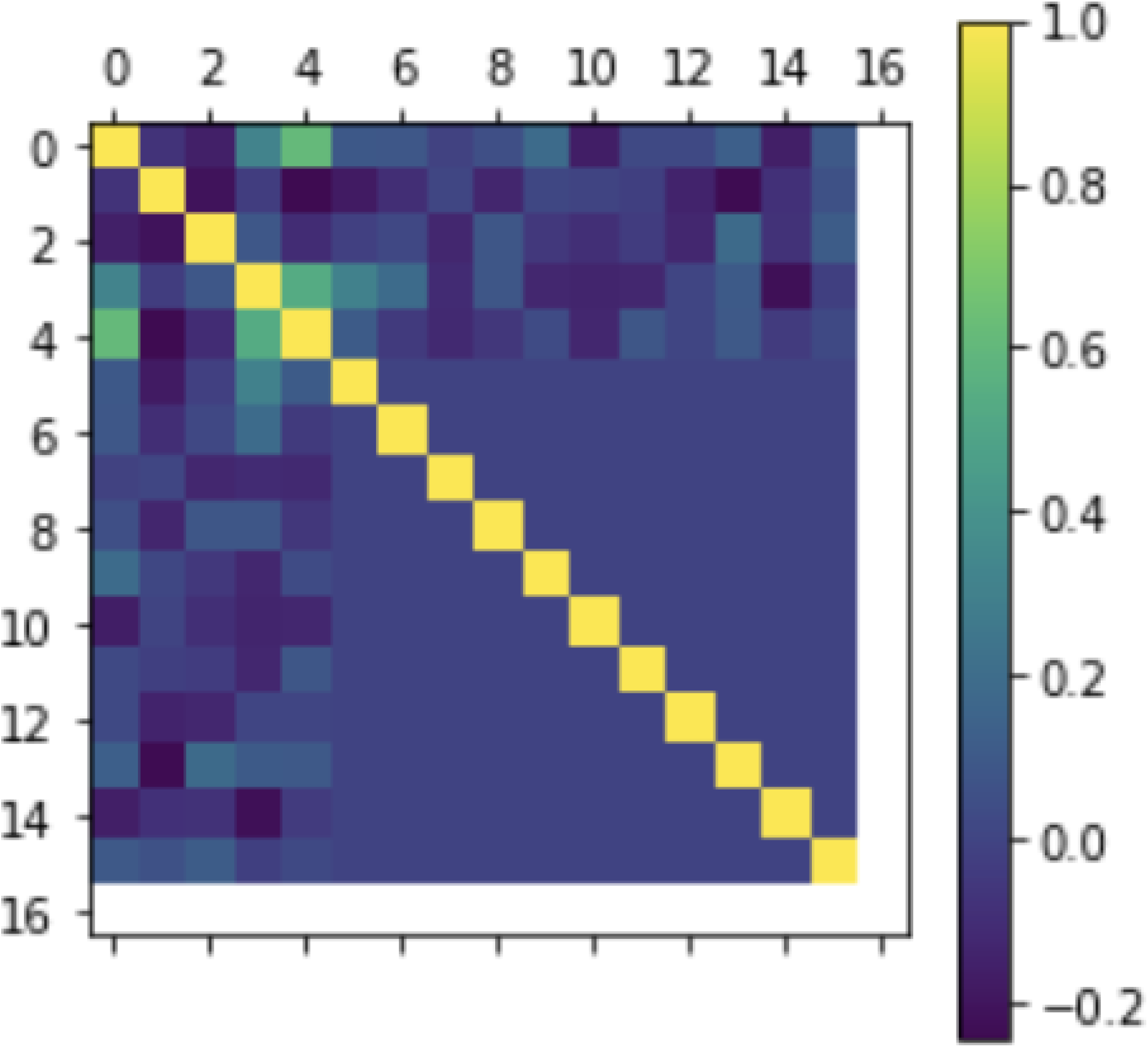
Pearson’s correlation coefficients between each predictor, the order of regressors are the same as shown in the design matrix in Figure 4

## Footnotes

1 The empirical distribution of references to story characters in pro-drop contexts is unbalanced as expected in naturalistic texts (See Section 2.6 for details). To help interpret accuracy levels, weighted logistic regression was applied in MVPA such that examples were weighted according to the prevalence of each class in the training data (This was realized by the scikit-learn (Pedregosa et al., 2011) class_weight= “balanced” option). The average accuracy from guessing randomly according to the empirical distribution in this weighted problem is 25%.

## Notes

### Competing Interest Statement

The authors have declared no competing interest.

